# The fungicide mancozeb induces astrocyte atrophy and disrupts Ca²⁺ signaling via inhibition of Orai1/STIM1-mediated SOCE

**DOI:** 10.64898/2026.06.12.731805

**Authors:** Ye-Ji Kim, Dong Ho Woo

**Author notes:** To whom correspondence should be addressed, Center for Global Biopharmaceutical Research, Korea Institute of Toxicology, Daejeon 34114, South Korea.

## Abstract

Mancozeb, a widely used fungicide composed of manganese ethylene-bis-dithiocarbamate with zinc salts, has raised concerns due to its potential neurotoxic effects. In this study, we investigated how chronic oral administration of mancozeb affects astrocyte function and neurobehavior in mice, focusing on store-operated Ca²⁺ entry (SOCE), mediated by Orai1 and STIM1. Mancozeb treatment at 0.5 µg/kg/day for 4 weeks reduced glial fibrillary acidic protein (GFAP) expression in the hippocampus and corpus callosum of mice, indicating astrocyte atrophy. Further, administration at the human acceptable daily intake (30 µg/kg/day) for 1 week induced hippocampal astrocyte atrophy and hyperlocomotor activity in open field tests. In vitro experiments revealed that mancozeb specifically inhibited SOCE in astrocytes by targeting the Orai1/STIM1 complex, as its inhibitory effect was abolished by short hairpin RNA (shRNA)-mediated knockdown of Orai1 or STIM1, but not by knockdown of TRPA1 or scramble shRNA. This demonstrates that mancozeb-mediated SOCE inhibition critically depends on the presence of Orai1 and STIM1, highlighting the molecular specificity of its action. Furthermore, mancozeb diminished endoplasmic reticulum (ER) Ca²⁺ stores and P2Y1 receptor agonist-induced Ca²⁺ transients. Electrophysiological analyses revealed that mancozeb selectively decreased the inhibitory postsynaptic current frequency without affecting excitatory currents, suggesting reduced astrocyte-mediated GABA release. Collectively, these findings demonstrate that mancozeb disrupts astrocytic Ca²⁺ homeostasis through Orai1/STIM1-dependent SOCE inhibition, leading to astrocyte atrophy and altered inhibitory neurotransmission, which may underlie the observed behavioral changes. These results highlight the potential neurotoxic risk posed by mancozeb via the impairment of astrocyte function and intracellular Ca²⁺ regulation. Importantly, these neurotoxic effects occurred at concentrations below current regulatory safety limits (ADI), indicating that mancozeb-induced disruption of astrocytic Ca²⁺ signaling provides a mechanistic basis for re-evaluating established human safety exposure standards.

**Environmental Implications:** Our findings highlight that the widespread use of mancozeb has a significant impact on brain health. Mancozeb was shown to induce astrocyte atrophy even at low concentrations, amounting to six times the human acceptable daily intake. Mancozeb causes impairment of GABAergic synaptic transmission of neurons by disrupting the Ca²⁺ homeostasis via inhibition of Orai1 and STIM1 of astrocytes. These findings indicate that current regulatory standards significantly underestimate the risks of long-term mancozeb exposure to brain health. Therefore, this study underscores the risks of astrocyte-mediated neurotoxicity resulting from pesticide residue ingestion and emphasizes the need to rigorously re-evaluate current exposure limits from the perspective of brain health.

## 1. Introduction

Manganese ethylene-bis-dithiocarbamate with zinc salts, commonly known as the fungicide mancozeb, was officially approved by the Environmental Protection Agency (EPA) for use in the growth of almonds, cabbage, lettuce, peppers, broccoli, walnuts, and tangerines in 2011 and 2013. However, the EPA has expressed concerns regarding the potential toxicity of mancozeb, not only in thyroid development and the reproductive system, but also in the nervous system [1]. Treatment of MES23.5, a rodent neuronal cell line, with mancozeb was shown to increase alpha-synuclein expression, suggesting that mancozeb can induce abnormalities in the central nervous system (CNS) [2]. Mancozeb has further been shown to exert neurotoxic effects in laboratory animals, including rats and mice. Studies have suggested that exposure to mancozeb can damage the nervous system, including the brain, spinal cord, and peripheral nerves. Specifically, mancozeb has been shown to trigger the degeneration of nerve fibers and disruption of nerve cell function [3], which can lead to a range of neurological symptoms, including tremors, coordination problems, and cognitive impairment, indicating that mancozeb causes neurodegenerative diseases such as Parkinson’s disease. One study suggested that exposure to mancozeb can negatively affect glial cells, including astrocytes [4]. Further, rats exposed to mancozeb showed alterations in the morphology and function of astrocytes associated with disruptions in neuronal signaling and increased susceptibility to neurotoxicity [5,6].

Astrocyte toxicity is generally explained by the upregulation of glial fibrillary acidic protein (GFAP). The physiological functions of astrocytes are closely related to their structurally intricate subcellular processes, as demonstrated by the visualization and quantification of GFAP, an astrocyte marker. Each individual astrocyte is in contact with hundreds of thousands of synapses [7]. Therefore, astrocyte architecture is critical for synaptic function in brain physiology. The tripartite synapse between these cells is composed of a presynaptic terminal and a postsynaptic membrane, including the microdomain and astrocyte membrane. Astrocytic Ca^2+^ release from the endoplasmic reticulum is critical for various physiological processes in the brain. Intracellular Ca^2+^ transients in astrocytes are important for the release of gliotransmitters, such as glutamate [8,9], GABA [10–12], D-serine [13–15], and ATP [16,17], which function to fine-tune synaptic activity. Astrocytes are local circuit modulators. The cellular mechanism involved in astrocytic morphology, which was identified by quantifying GFAP staining. Upregulation of GFAP and hypertrophy are known mediators of inflammation; however, the effects of the downregulation of GFAP, hypotrophy, or atrophy are not well characterized.

The transient expression of nuclear factor IA (NFIA) in pluripotent stem cells increases the number of functional astrocytes and GFAP-positive cells, indicating that NFIA expression is correlated with GFAP expression [18]. NFIA is required for the long-term inhibition of neurogenesis, suggesting that it is closely related to gliogenesis [19]. Conditional knockout mice with NFIA showed reduced GFAP expression, while the selective ablation of NFIA failed to produce reactive astrocytes (quantified by counting GFAP-expressing cells), suggesting that NFIA is a promoter of the gfap gene [20]. Conditional knockout of NFIA results in hippocampal and cortical-specific GFAP atrophy [21], suggesting a correlation between NFIA and GFAP expression.

The calcium release-activated calcium channel protein 1(Orai1)-Stromal Interaction Molecule 1 (STIM1) complex contributes to Ca^2+^ homeostasis in astrocytes. Once Ca^2+^ is depleted in the endoplasmic reticulum (ER), store-operated Ca^2+^ entry (SOCE) occurs via the interaction of Orai1 in the cytoplasmic membrane and STIM1 in the ER membrane. SOCE replenishes the ER Ca^2+^ reserves. In the brain, astrocytes facilitate ER Ca^2+^ to use Ca^2+^ transients to form Ca^2+^ oscillations and modulate neuronal activity [22] [23], gliotransmitters [24–26], gene expression, and vascular functions [27]. SOCE homeostasis can influence the shape of glial cells and modify local signaling. Herein, we investigated the molecular mechanisms underlying astrocyte atrophy in SOCE.

## 2. Materials and methods

### 2.1. Animals

Postnatal day (P) 1 Sprague Dawley (SD) rats and mice for the primary cultured neurons were purchased (Orient Bio, Korea). All experiments were conducted in accordance with animal study protocols approved by the Institutional Animal Care and Use Committee of the Korea Institute of Toxicology (IAC-23-01-0335-0184, IAC-23-01-0383-0166, and IAC-23-01-0564-0325, KIT, Daejeon, Korea). Animal care was performed following National Institutes of Health (NIH) guidelines. 5 weeks male mice were introduced into our animal facility and acclimated for 1 week. For 4 weeks, oral administration of 0.5 μg/kg mancozeb was administered from every Monday to Friday. For 1 week, oral administration (acceptable daily intake, ADI) of 0.023 μg/kg was orally administered every day. All mice were anesthetized with 20 mg/ml/kg Avertin and perfused with 4% paraformaldehyde (PFA, J19943, Thermo Fisher Scientific, MA, USA) for fixation.

### 2.2. Immunohistochemistry

For 4-weeks experiment (Figure 1, 0.5 μg/kg/day) and 1-week experiment (Figure 2, 0.023μg/kg/day), the fixed brain was frozen with OCT compound and sliced with a 20 μm thickness. Brain slices were washed with Phosphate-Buffered Saline (PBS, LB001-02, Welgene, Gyeongsan, Korea) 3 times and incubated with a blocking solution for 1 h (0.5 % Triton X-100 (X-100, Sigma, MA, USA), 1 % BSA (BSAS 0.1, Bovogen, Melbourne, Australia), 5 % normal goat serum (S-1000, Vector Laboratories, Inc., CA, USA) in PBS) at room temperature (RT). Primary antibodies in the blocking solution were incubated overnight at 4°C. Anti-GFAP (13-0300, 1:500 dilution, Invitrogen, MA, USA), Anti-NFIA (ab228897, 1:200, Abcam, CB, UK) were used as primary antibodies. After washing with PBSTB (0.5 % Triton X-100, 1 % BSA in PBS) 2 times for 10 minutes, cells were incubated with secondary antibodies in PBSTB for 1 h at RT. Alexa Fluor 488 goat anti-rat (A11006, 1:500, Invitrogen, CA, USA) and Alexa Fluor 594 goat anti-rabbit (A11012, 1:500, Invitrogen, CA, USA) were used as secondary antibodies. And washing with PBST for 10 minutes 2 times and PBS for 10 minutes 1 time at RT, brain slices on coverslips were incubated with a mounting medium including DAPI (H-1200-10, VECTASHIELD, CA, USA) and mounted onto a slide glass. A series of fluorescent images was obtained by a confocal microscope (Olympus, Tokyo, Japan) and analyzed by Image J software.

**Fig. 1.**
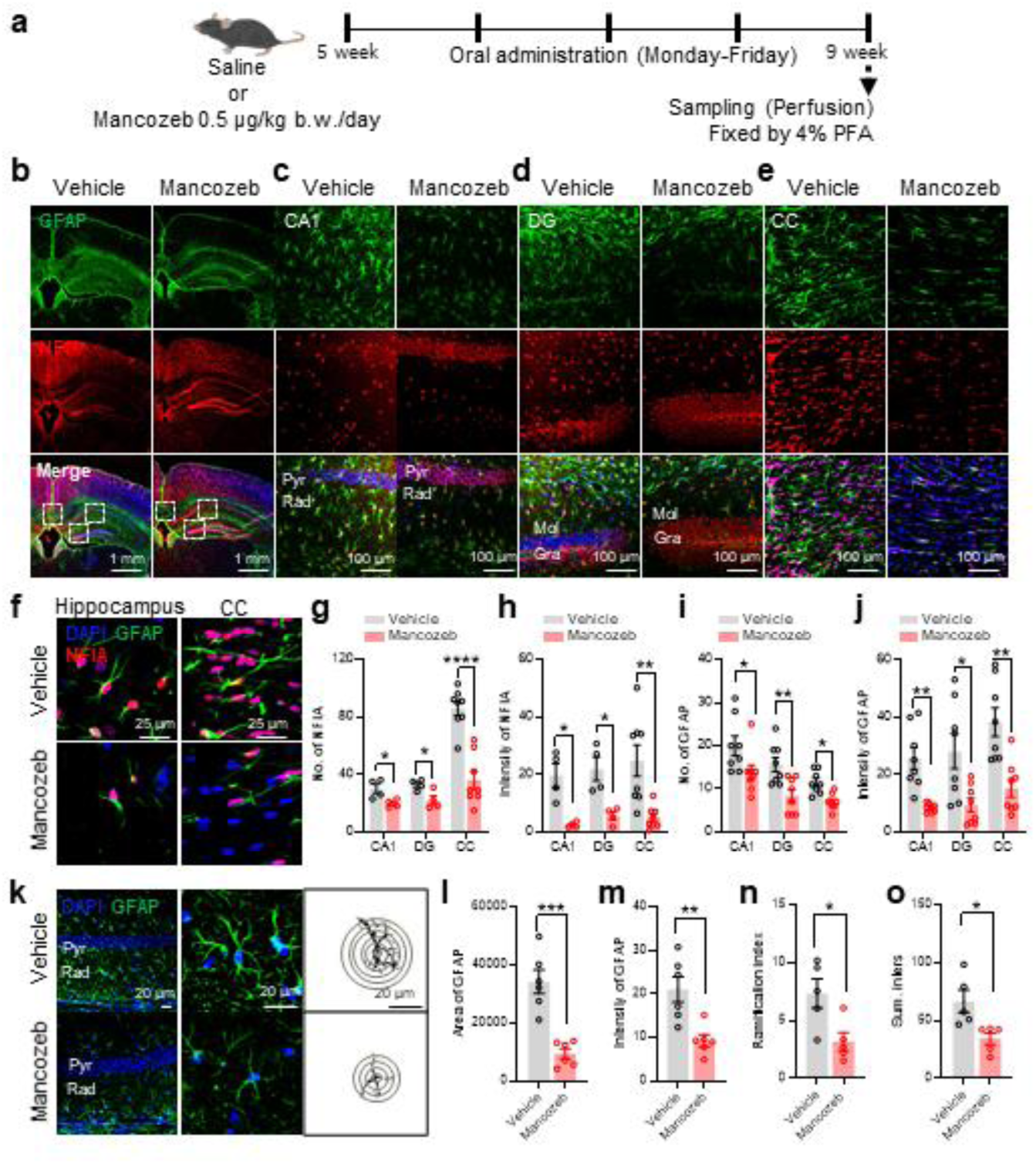
Orally low-dose intoxication of 0.5 μg/kg/day mancozeb is accompanied by atrophy of mouse hippocampal astrocytes including corpus callosum. **a-e** Experimental timeline for 4weeks-oral administration **(a)**. Coronal section of the mouse for low magnification including the hippocampus and the corpus callosum with glial fibrillary acidic protein (GFAP), Nuclear Factor I A (NFIA), and merge images without and with mancozeb. White dot boxes in merged images as corpus callosum, granular and molecular layer of the dentate gyrus (DG), and stratum radiatum (Rad) of hippocampal CA1 **(b)**. Scale bar 1mm. High magnification of hippocampal CA1 **(c)**, DG **(d)**, CC **(e)** without and with mancozeb. Pyramidal layer (Pyr), Stratum radiatum (Rad), Molecular layer (Mol), and granular layer (Gra). Scale bar 100 μm. **f-j** High magnification for DAPI, GFAP and NFIA of hippocampal rad and cc without and with mancozeb **(f)**. Scale bar 25 μm. Summary bar graph for number of NFIA **(g)**, intensity of NFIA **(h)**, number of GFAP **(i)**, and intensity of GFAP **(j)**. **k-o** High magnification of DAPI and GFAP from hippocampal rad and Sholl analysis for GFAP area without and with mancozeb **(k)**. Summary bar graph for area of GFAP **(l)**, intensity of GFAP **(m)**, ramification index **(n)**, and summed intersection (Sum. Inters, **o**). Pyramidal layer (Pyr), Stratum radiatum (Rad). Scale bar 20 μm. Statistical significance was evaluated with two-tailed unpaired Student’s t-test (*p < 0.05, **p < 0.01, ***p < 0.001, ****p < 0.0001). Data are presented as mean ±SEM.

**Fig. 2.**
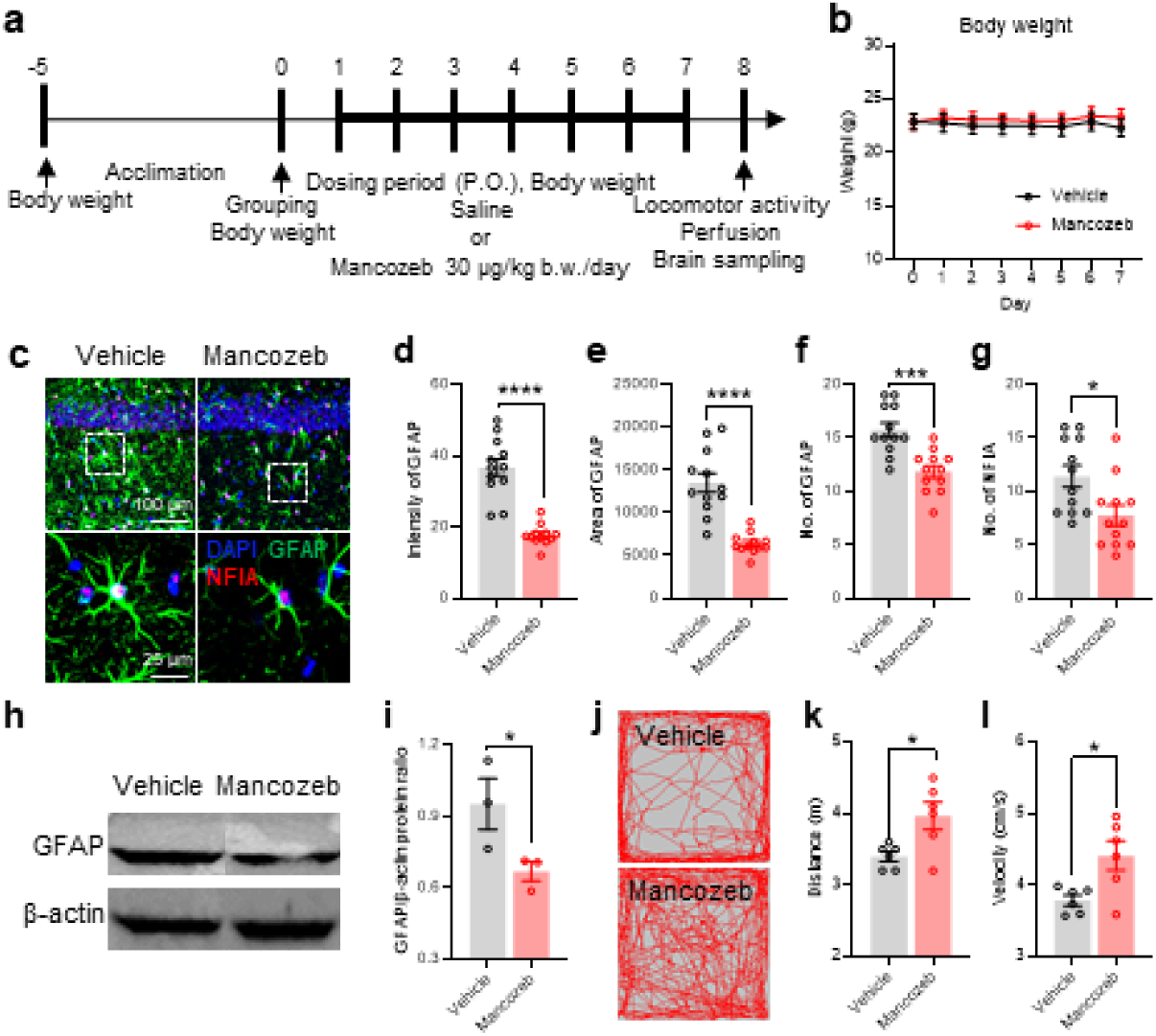
Orally ADI-dose intoxication of 30 μg/kg/day mancozeb shows atrophy of mouse hippocampal astrocytes and increases the distance of locomotor activity. **a** Experimental timeline for consecutive 7 days. **b** Line plots for the change of body weight along experimental procedure. **c-g** High magnification of DAPI, GFAP and NFIA of hippocampal rad with and without mancozeb **(c)**. Summary bar graph for intensity of GFAP **(d)**, area of GFAP **(e)**, number of GFAP **(f)**, and number of NFIA **(g)**. **h, i** Western blot for GFAP quantification without and with mancozeb **(h)** and summary bar graph for normalized GFAP **(i)**. **j-l** Cumulative trajectories of locomotion activity of the mouse **(j)**. Summary bar graph for cumulative moving distance **(k)** and velocity **(l)**. Statistical significance was evaluated with two-tailed unpaired Student’s t-test (*p < 0.05, ***p < 0.001, ****p < 0.0001). Data are presented as mean ±SEM.

### 2.3. Sholl analysis

Sholl analysis was performed using ImageJ software. Images obtained using a confocal microscope were acquired, and the soma of each astrocyte was identified as the center point. A series of circles (with 10 µm interval) was overlaid on the image. The number of times the processes intersected each circle and the ramification index were counted using automated software.

### 2.4. Western blotting

3 pup’s brains were seeded to a 60 mm dish with the protocol of primary astrocyte culture. 10 μM Mancozeb was treated for 24 hours at 70% confluency. The astrocytes were harvested and washed with cold phosphate-buffered saline (PBS, CBP007B, LPS, South Korea). Then the washed astrocytes were changed into RIPA lysis buffer with protease inhibitor cocktail for 20 min on ice. Lysis was centrifuged at 12000rpm at 4°C. The supernatant was quantified with the BCA assay (Thermo, MA, USA). 20 μg/sample were prepared with 4X sample loading buffer (BioRad, CA, USA) with 20 μl volume and loaded into the lane of the stacking gel. 100V for 1.5 hours was running on the running buffer of SDS-PAGE tank (BioRad, CA, USA) [28]. The gel was transferred to a PVDF membrane into a gel transfer at 20V for overnight at 4°C. The PVDF membrane was washed with PBS and incubated for 1 hour with a blocking solution (PBS, 0.05% tween 20, 5% skim milk). The membrane was treated with the primary antibody against GFAP for overnight at 4°C.

### 2.5. Open fields test

1 week-orally administered mice were first acclimated for one week in polycarbonate cages with bedding after acquisition before assessing their activity. The mice were grouped based on body weight. The mice were group-housed under a 12-h light/dark cycle at 25℃ with free access to food and water. The box (42×42 cm) used for the OFT was equipped with a recording camera. Each mouse was placed in the center of the box, habituated for 30 minutes, and then tested for 20 minutes. The tests were videotaped, and the distances traveled by the mice were analyzed using EthoVisionXT software (version 14, Noldus, Netherlands).

### 2.6. Primary rat and mouse cortical astrocyte culture

Rat primary cortical astrocytes were prepared from P1 of SD rats as described [29]. The cerebral cortex of the rat was dissected free of adherent meninges, minced, and dissociated into a single-cell suspension by trituration. Astrocytes were grown in Dulbecco’s modified Eagle’s medium (DMEM, Gibco, MA, USA) supplemented with 10% heat-inactivated horse serum, 10 % heat-inactivated fetal bovine serum (FBS, Gibco, MA, USA), and 1% penicillin-streptomycin (100 units/ml penicillin–0.1 mg/ml streptomycin, Gibco, MA, USA). Cultures were maintained at 37°C in a humidified 5% CO_2_ incubator. On the days in vitro (DIV) 3, cells were vigorously washed with repeated pipetting, and the media was replaced to get rid of debris and other floating cell types. On DIV 7-8, cells were replated onto a coverglass (8×10^4^ per well) coated with 50 µg/ml Poly-D-Lysine (PDL, Merck, USA) for imaging experiments.

### 2.7. Calcium imaging

The external solution contained (mM): 150 NaCl, 10 HEPES, 3 KCl, 2 CaCl2, 2 MgCl2, 5.5 glucose; pH adjusted to pH 7.3 - 7.4 and osmolarity to 325 - 330 mOsm/kg. Intensity images of 510 nm wavelength were acquired from excitation wavelengths at 340 and 380 nm using an Electron Multiplying Charge-Coupled Device (EMCCD) camera (Ixon 867, Andor Oxford instrument). The neurons were incubated for 40 minutes with 5 μM fura-2 AM (Thermofisher, MA, USA) and 5 μl pluronic F-127 (Thermofisher, MA, USA). Images are acquired every 1 sec. In the field of view, 50 cells were measured and analyzed.

For SOCE, 1 μM thapsigargin for 2 min (TG) releases Ca^2+^ from astrocytic ER to deplete the Ca^2+^ store of ER, called Ca^2+^ release, and 2 mM extracellular Ca^2+^ was applied for the last to induce SOCE, called Ca^2+^ entry. For the treatment of SOCE inhibitor, mancozeb, SKF96365 for Orai1 channel inhibitor, and HC030031 for TRPA1 inhibitor were pretreated for 3 min prior to 2 mM Ca^2+^ application (Fig. 3).

**Fig. 3.**
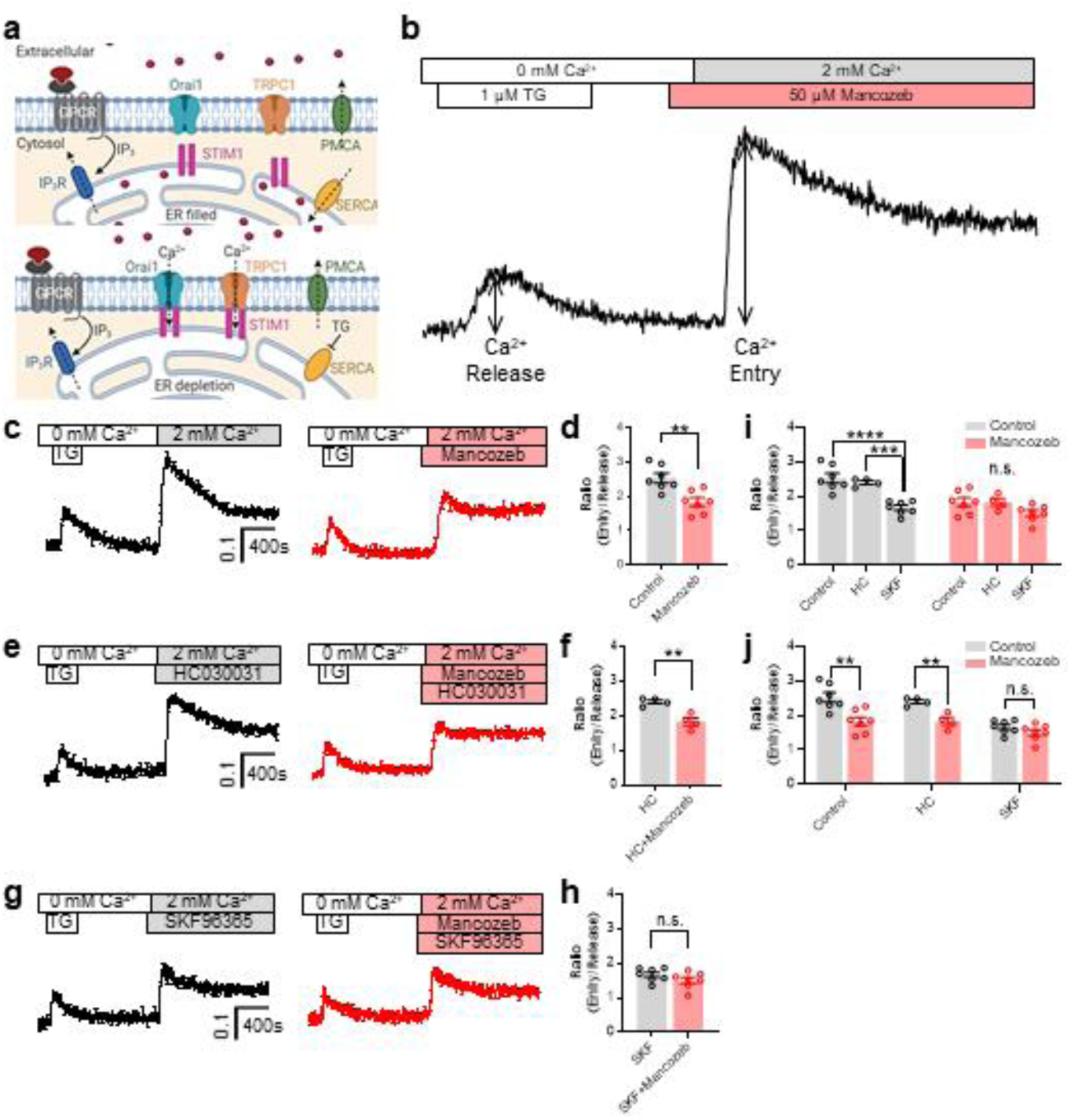
Pharmacological evidence of involvement of mancozeb in store-operated Ca^2+^ entry (SOCE). **a** Schematic cartoon for indicated drug targets before (upper panel) and after (lower panel) Ca^2+^ depletion of endoplasmic reticulum (ER). **b** An experimental procedure and the representative trace of SOCE. **c** Representative traces of SOCE and the summary bar graph for the evidence that mancozeb targets SOCE. Thapsigargin (Tg)-depleted ER Ca^2+^ (Ca^2+^ release) and 2 mM extracellular Ca^2+^-induced Ca^2+^ entry (left black) was blocked by mancozeb (right red). **d** Summary bar graph for the ratio (Ca^2+^ entry / Ca^2+^ release). **e** Tg-depleted ER Ca^2+^ (Ca^2+^ release) and 2 mM extracellular Ca^2+^-induced Ca^2+^ entry with HC-030031, TRPC channel inhibitor (left black) were blocked by mancozeb (right red). **f** Summary bar graph for the ratio. **g** Tg-depleted ER Ca^2+^ (Ca^2+^ release) and 2 mM extracellular Ca^2+^-induced Ca^2+^ entry with SKF96365, Orai1 channel inhibitor (left black) were not blocked by mancozeb (right red). **h** Summary bar graph for the ratio. **i** Summary bar graph for showing that the main mechanism of SOCE is Orai1. **j** Summary bar graph for showing that mancozeb targets Orai1. Statistical significance was evaluated with two-tailed unpaired Student’s t-test (**d, e, h,** and **j,** **p < 0.01) and two-way ANOVA followed by Tukey’s test for multiple comparisons (**i,** ***p < 0.001, ****p < 0.0001). Data are presented as mean ±SEM.

For monitoring receptor-mediated Ca^2+^ transients, astrocytes were treated with 0.5, 5, and 25 μM mancozeb for 24 hours. On an experimental day, 1 μM MRS2365, a specific purinergic receptor P2Y1 agonist (Fig. 5).

### 2.8. Immunocytochemistry

Cultured astrocytes on the coverslip were fixed for 15 min in 4% paraformaldehyde (PFA, J19943, Thermo Fisher Scientific, MA, USA). Astrocytes were washed with PBS (LB001-02, Welgene, Gyeongsan, Korea) 3 times and incubated with a blocking solution for 1 h at RT. Primary antibodies in the blocking solution were incubated overnight at 4°C. Anti-GFAP (13-0300, 1:1000 dilution, Invitrogen, MA, USA) was used as a primary antibody. After washing with PBSTB 2 times for 10 minutes, cells were incubated with secondary antibodies in PBSTB for 1 h at RT. Alexa Fluor 488 goat anti-rat (A11006, 1:500, Invitrogen, CA, USA) was used as a secondary antibody. And washing with PBST for 10 minutes 2 times and PBS for 10 minutes 1 time at RT, brain slices on coverslips were incubated with a mounting medium including DAPI (H-1200-10, VECTASHIELD, CA, USA) and mounted onto a slide glass. A series of fluorescent images was obtained by a confocal microscope (Olympus, Tokyo, Japan) and analyzed by Image J software.

### 2.9. Electroporation for gene silencing

shRNA vectors were used to specifically knock down the expression of target genes. shRNAs targeting Orai1 (shRNA of Orai1), TRPA1 (shRNA of TRPA1), TRPC1 (shRNA of TRPC1), and STIM1 (shRNA of STIM1) were designed to selectively bind to their respective mRNAs as described below.

Orai1, ‘5′-GCACCTGTTTGCCCTCATGAT-3′’;

Trpa1: 5′- GCAAGCTTCCTTTCTGCATAT-3′ [30]

Trpc1, ‘5′-CTATGGCTTAGCTACTTTGA-3′’;

Stim1, ‘5′-GCGAGATGAGATCAACCTTGC-3′’

pENSR-shRNAs for Orai1, Orai2, Orai3, Trpc1, and Stim1 were constructed by using the EZchangeTM site-directed mutagenesis kit (Enzynomics) and confirmed by sequencing [31]. Electroporation of shRNA was performed using an electroporator (Invitrogen Neon transfection system, CA, USA). Astrocytes were harvested and resuspended in 100 μl Neon Resuspension Buffer (1.2 × 10^6^ cells). Cell suspension and each shRNA plasmid were gently mixed (1 μg per 1.2 × 10^6^ cells). The mixture was aspirated into a Neon pipette tip (100 μl) and subjected to electroporation using optimized parameters (1300V, 20 ms, 2 pulses). After electroporation, cells were immediately transferred to pre-warmed culture medium (DMEM with 10% FBS) onto PDL-coated coverglass in a 24-well plate and incubated at 37°C with 5% CO₂ for recovery.

### 2.10. Slice preparation

Mice were anesthetized with isoflurane and perfused with ice-cold oxygenated (95% O2, 5%CO2) artificial cerebrospinal solution (ACSF) composed of (mM): 130 NaCl, 1.25 NaH_2_PO_4_, 3.5 KCl, 24 NaHCO_3_, 1.5 CaCl_2_, 1.5 MgCl_2_, and 10 glucose (pH 7.4 by NaOH; osmolarity adjusted to 315–320 mOsm/kg). The brain was dissected and mounted on a vibratome (Leica1200s, Wetzlar, Germany). Horizontal slices (300 μm thick) were prepared using the vibratome and stored in an incubation solution (ACSF) at room temperature.

### 2.11. Whole-cell patch-clamp recording

Whole-cell patch recording from the hippocampal CA1 region of the brain slice under voltage clamp (holding potential –70 mV) was made with a MultiClamp 700B amplifier digitized by Digidata 1322A data acquisition system (Molecular Devices, CA, USA). The recording chamber was continually perfused with recording solution composed of (mM): 130 NaCl, 1.25 NaH_2_PO_4_, 3.5 KCl, 24 NaHCO_3_, 1.5 CaCl_2_, 1.5 MgCl_2_, and 10 glucose (pH 7.4 by NaOH; osmolarity adjusted to 315–320 mOsm/kg). Recording electrodes (4 – 7 MΩ) for sEPSC were filled with (mm): 150 CsMeSO_4_, 10 NaCl, 0.5 CaCl_2_, 10 Hepes, (pH adjusted to 7.3 with CsOH and osmolarity adjusted to 310 mOsm/kg). Recording electrodes (4 – 7 MΩ) for sIPSC were filled with (mm): 150 KCl, 10 NaCl, 0.5 CaCl_2_, 10 Hepes, (pH adjusted to 7.3 with CsOH and osmolarity adjusted to 310 mOsm/kg). For inhibition of the voltage-gated sodium channel, 0.5 mM QX 314 bromide (1014, Tocris, Bristol, US) was added to the internal solution. All electrophysiological data from cultured cells in this study were recorded using temperature controller (23–25◦C, Axon Instrument, CA, USA). Mini Analysis (Synaptosoft, NJ, USA) was used to analyze frequency and amplitude. GraphPad Prism (version 10) was used for one-way ANOVA post hoc test.

### 2.12. Statistical analysis

The numbers and individual dots refer to the number of cells or animals unless otherwise clarified in the figure legends. GraphPad Prism 9.1.2 (GraphPad Software) was used for data presentation and statistical analysis. For the electrophysiological experiment, Mini analysis (Synaptosoft) and Clampfit (Molecular Devices, MA, USA) were used. For plotting electrophysiological data, the Sigma plot (Systat, CA, USA) was used. Statistical significance was set at *p < 0.05, **p < 0.01, ***p < 0.001, ****p < 0.0001. Data are presented as mean ± SEM.

## 3. Results

### 3.1. Characterization of astrocyte atrophy in the hippocampus, including the corpus callosum, in mice following intoxication of orally low-dose mancozeb for 4 weeks

We planned to feed one-sixtieth of the ADI amount of mancozeb for one month, anticipating that this regimen would induce brain inflammation accompanied by hypertrophy of the astrocyte marker GFAP. The ADI is the amount of mancozeb ingested by individuals who consume apples, broccoli, leeks, and cauliflowers daily, calculated as a simulation of the amount remaining when ADI is applied to one apple and stored for 30 days following the washing process [32]. We orally administered 0.5 μg/kg/day mancozeb for 4 weeks to determine the low concentration and chronic effect of mancozeb (Fig. 1a).

Weekly monitoring by rough observation revealed some behaviors that appeared unnatural and persisted for 4 weeks. Surprisingly, GFAP atrophy was observed in a variety of brain regions, including the stratum radiatum (Rad) of the hippocampal CA1, hilus of the dentate gyrus (DG), and corpus callosum (Fig. 1b-e). Since NFIA is strongly related to GFAP atrophy, we measured the intensity and area of fluorescence in NFIA-positive cells [18–21]. Mancozeb significantly decreased the fluorescence intensity and number of NFIA and GFAP cells in the CA1, DG, and CC (Fig. 1f-j). In addition to the intensity and number of fluorescences, mancozeb significantly increased the index of GFAP atrophy, area, intensity, ramification, and sum of the intersection in CA1 (Fig. 1k-o).

### 3.2. Induction of atrophy in hippocampal CA1 astrocytes in mice following intoxication of ADI-dose mancozeb administered orally for 1 week

We subsequently investigated whether continuous oral administration of ADI for 7 days would induce atrophy of GFAP. To achieve this, 30 μg/kg/day mancozeb was orally administered to mice for 7 consecutive days. This treatment showed no effects on body weight (Fig. 2a-b), suggesting that mancozeb does not exert severe systemic toxicity. However, similar to chronic mancozeb oral administration for 4 weeks, atrophy was also induced by subchronic mancozeb administration for 7 days (Fig. 2c).

The intensity (Fig. 2d), area (Fig. 2e), and number (Fig. 2f) of GFAP fluorescence signals were reduced after 7 days of oral administration of ADI mancozeb. NFIA staining also decreased (Fig. 2g). These results were similar to the atrophy of GFAP and NFIA following the chronic administration of mancozeb for 4 weeks. Western blotting showed a reduction in GFAP expression normalized with β–actin (a constitutively-expressed protein used for protein expression normalization) (Fig. 2h, i). Before the mice were euthanized, an open field test (OFT) was conducted to assess general behavior. Interestingly, mancozeb increased locomotion (Fig. 2j, k) and movement speed (Fig. 2j, l), indicating that GFAP atrophy is highly related to hyperlocomotor activity. These results suggest that the hyperlocomotor activity induced by mancozeb ingestion is similar to the withdrawal phenomenon of drug addiction.

### 3.3. The Orai I/STIM1 complex is the molecular target of mancozeb, identified by pharmacology and gene silencing

Calcium imaging was performed to confirm that mancozeb inhibited SOCE. Orai1/STIM1 complex is a major channel component for SOCE [33,34]. When endoplasmic reticulum (ER) Ca^2+^ levels are depleted, the entry of external calcium occurs via the Orai1/STIM1 complex (Fig. 3a). ER Ca^2+^ depletion was induced by treatment with 1 μM thapsigargin (TG), an inhibitor of the H^+^ ATPase, which pumps Ca^2+^ into the ER in the absence of external Ca^2+^. Once the ER Ca^2+^ was depleted, external 2 mM Ca^2+^ was added as Ca^2+^ entry with 50 µM mancozeb treatment, including pretreatment for 2 minutes (Fig. 3b). The ratio of Ca^2+^ entry to Ca^2+^ release was calculated. Mancozeb reduced this ratio and normalized Ca^2+^ entry (Fig. 3c, d, Tables 1 and 2), indicating that mancozeb inhibited SOCE. HC030031, a TRPA1 inhibitor, blocked this mancozeb-reduced SOCE (Fig. 3e, f, Tables 1 and 2), indicating that mancozeb inhibition does not share the same pathway as HC030031. SKF96365, an Orai1 inhibitor, did not block mancozeb-reduced SOCE (Fig. 3g, h, Tables 1 and 2), suggesting that mancozeb-inhibited Ca^2+^ entry is related to the pathway involving Orai1. Normalized Ca^2+^ entry with HC030031 and SKF96365 was compared to that without mancozeb (Fig. 3i, Tables 1 and 2), indicating that Orai1 is a major mechanism for SOCE. Mancozeb did not decrease the normalized Ca^2+^ entry in the presence of SKF96365 (Fig. 3j, Tables 1 and 2), indicating that mancozeb shares the Orai1 pathway.

**Table 1.** Calcium transients analysis to measure changes in SOCE.

| No Mancozeb | Release | Entry | Ratio |
| --- | --- | --- | --- |
| Control | 0.098±0.006 | 0.247±0.017 | 2.531±0.138 |
| HC030031 | 0.113±0.012 | 0.272±0.032 | 2.408±0.058 |
| SKF96365 | 0.098±0.007 | 0.162±0.006 | 1.671±0.076 |

**Table 2.** Calcium transients analysis to measure changes in SOCE with mancozeb treatment.

| Mancozeb treatment | Release | Entry | Ratio |
| --- | --- | --- | --- |
| Control | 0.103±0.010 | 0.185±0.019 | 1.856±0.125 |
| HC030031 | 0.121±0.013 | 0.219±0.015 | 1.830±0.111 |
| SKF96365 | 0.093±0.013 | 0.137±0.013 | 1.506±0.151 |

To increase our confidence in the molecular mechanisms of mancozeb, we used gene silencing to determine whether it targets the Orai1/STIM1 complex. Mancozeb reduced Ca^2+^ entry into rat primary astrocytes transfected with scrambled shRNA (Fig. 4a-c), Trpa1 shRNA (Fig. 4d, e), and Trpc1 shRNA (Fig. 4f, g), suggesting that mancozeb decreased SOCE by inhibiting the Orai1 channel. However, mancozeb did not decrease Ca^2+^ entry from astrocytes expressing Orai1 shRNA (Fig. 4h, i), and Stim1 shRNA (Fig. 4j, k), suggesting that mancozeb targets Orai1/Stim1. Orai1/Stim1 is a pivotal factor in SOCE (Fig. 4l), and mancozeb did not reduce Ca^2+^ entry from astrocytes expressing Orai1 and Stim1 shRNA (Fig. 4m), suggesting that mancozeb targets Orai1 and Stim1.

**Fig. 4.**
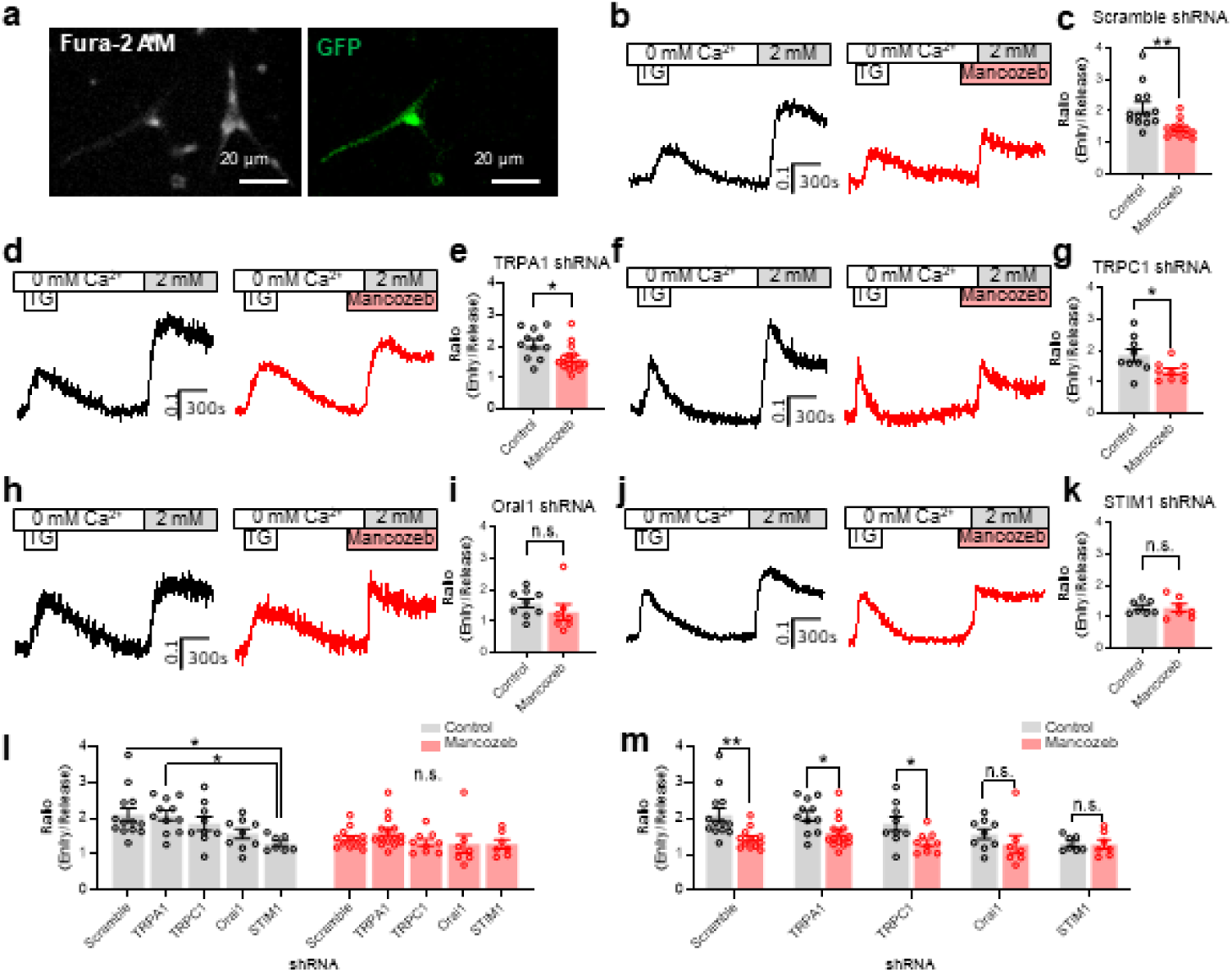
Mancozeb targets the Orai1-STIM1 complex. **a** Primary cultured astrocytes stained with the fura-2 image and astrocytes expressing GFP. **b** Representative traces for Ca^2+^ release and Ca^2+^ entry without (left black trace) and with mancozeb (right red trace) from mouse cortical primary cultured astrocytes expressing shRNA of scramble. **c** Summary bar graph for ratio (entry/release). **d, f, h, and j** Representative traces for Ca^2+^ release and Ca^2+^ entry without (left black trace) and with mancozeb (right red trace) from mouse cortical primary cultured astrocytes expressing shRNA of TRPA1 **(d)**, TRPC1 **(f)**, Orai1 **(h)**, and STIM1 **(j)**. **e, g, i, and k** Summary bar graph for the ratio. **l** Summary bar graph showing that Orai1 and STIM1 are targets for SOCE. **m** Summary bar graph showing that mancozeb targets Orai1 and STIM1. Statistical significance was evaluated with two-tailed unpaired Student’s t-test (**c, e, g, I, k, and m,** *p < 0.05, **p < 0.01) and two-way ANOVA followed by Sidak’s test for multiple comparisons (**l,** *p < 0.05). Data are presented as mean ±SEM.

**Fig. 5.**
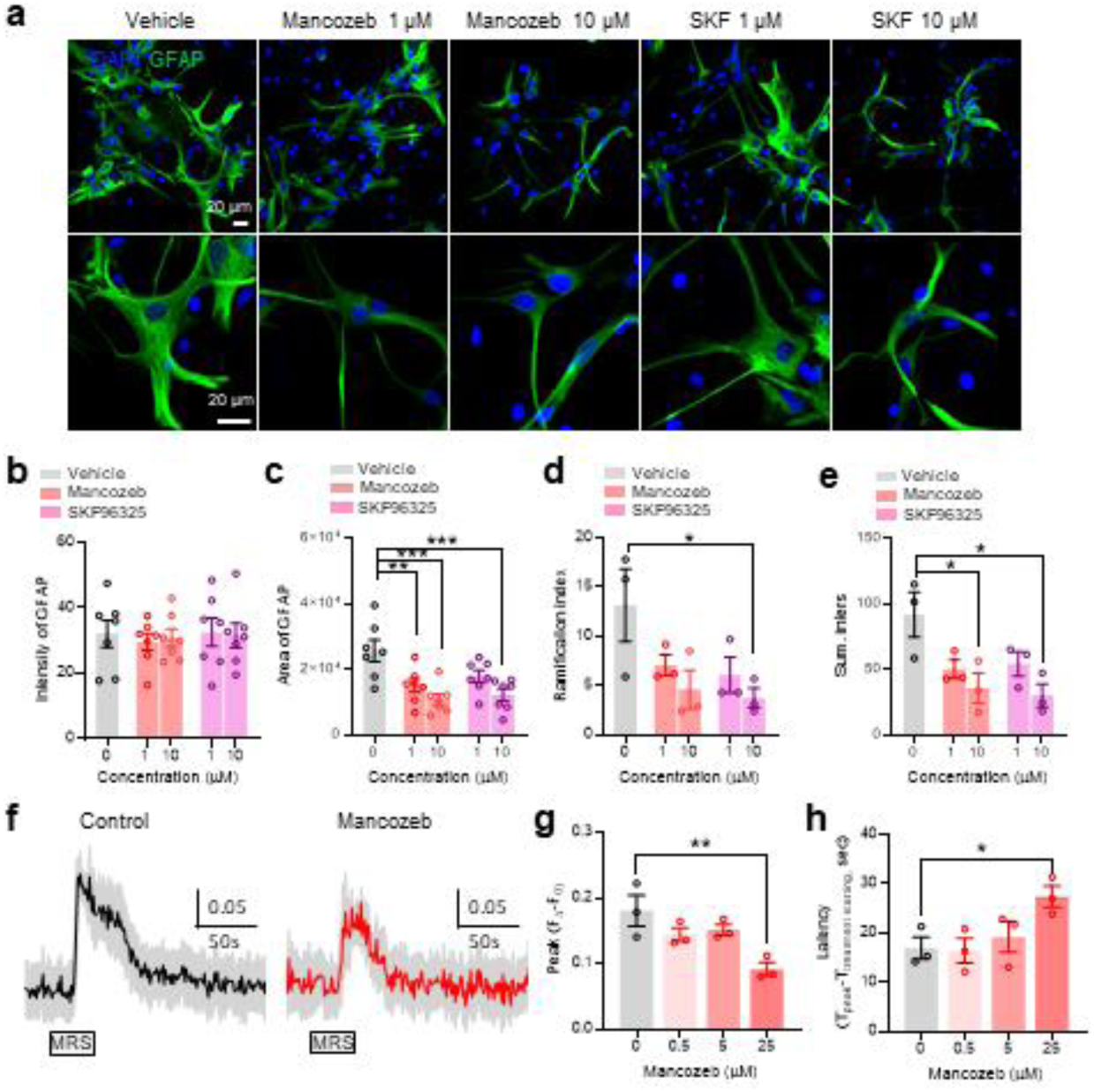
SKF96365, Orai1-STIM1 inhibitor, shows atrophy of GFAP. **a** Low and high magnification of GFAP fluorescence images without and with 1 μM mancozeb, 10 μM mancozeb, 1 μM SKF96325, and 10 μM SKF96325 after 24 hours of incubation. Scale bar 20 μm. **b-e** Summary for intensity of GFAP **(b)**, area of GFAP **(c)**, ramification index of GFAP **(d)**, and summed intersections of GFAP **(e)**. **f** Representative trace following activation by MRS2365, a selective P2Y1 receptor agonist. **g, h** Summary bar graph for peak amplitude **(g)** and latency **(h)**. Statistical significance was evaluated with one-way ANOVA followed by Dunnett’s test for multiple comparisons (*p < 0.05, **p < 0.01, ***p < 0.001). Data are presented as mean ±SEM.

### 3.4. Implications of SOCE on astrocyte atrophy

Prior research has shown that when murine epidermal cells were treated with benzohydroquinone, an Orai1 activator, cell thickness and proliferation increased in a manner dependent on the Orai1 inhibitor [35]. In addition, astrocytic Ca^2+^ is known to be a key molecule in the structural reorganization of plasticity in the connectome of the central nervous system via morphological changes [36]. Astrocytic Ca^2+^ controls the release of substances that are important for synaptic modulation and pathological conditions [9,11,17,25,37], while astrocytic Ca^2+^ signaling is closely related to astrocytic structures [38]. The structure of the ER and mitochondria can change astrocyte structure [39]. Therefore, we speculated that astrocytic ER Ca^2+^ influences astrocyte architecture. SOCE is a well-known mechanism of ER Ca^2+^ replenishment via interaction of the cytoplasmic membrane Orai1 channel and ER membrane STIM1 [33,40,41].

Given the above research, we aimed to investigate whether glial cell structure can be altered by SOCE. SKF96365, an Orai1 channel inhibitor, was assessed to see whether it induced atrophy to the same extent as mancozeb treatment. Mancozeb reduced the area of GFAP to the same extent as SKF963659, but did not decrease the intensity of GFAP (Fig. 5a-c). Sholl analysis showed a reduced ramification index and summed intersection (Fig. 5d, e), indicating that mancozeb-mediated morphological changes were very similar to those of SKF96365, an Orai1 channel inhibitor. We surmised that mancozeb may reduce the Ca^2+^ levels in the ER by inhibiting SOCE. MRS2365, a selective P2Y1 agonist, was used to induce the receptor-mediated Ca^2+^ transients. We expected these receptor agonists to reduce the peak and latency of Ca^2+^ transients. Mancozeb reduced the peak amplitude and delayed the latency of the MRS2365-induced Ca^2+^ transients (Fig. 5f-h).

### 3.5. Recovery of mancozeb-induced sIPSC reduction via Ca^2+^ coadministration ^+^

Previously, we showed that mancozeb reduces glial calcium transients through the inhibition of SOCE in glial cells, thereby depleting endoplasmic reticulum Ca^2+^ stores. In the previous study shows that mutant mice with 1,4,5-trisphosphate receptors (IP3R2), a major component of ER Ca^2+^ signaling, exhibited not only a decrease in Ca^2+^ but also a consequent decrease in GABAergic neurotransmission [42]. Therefore, our subsequent investigation focused on whether mancozeb treatment influences the release of the neurotransmitters, GABA and glutamate. The release of GABA and glutamate was recorded in postsynaptic CA1 pyramidal neurons in the hippocampus (Fig. 6a). Under acute mancozeb treatment, while spontaneous inhibitory postsynaptic current (sIPSC) frequency reduced (Fig. 6b-d), spontaneous excitatory postsynaptic current (sEPSC) frequency was unaffected (Fig. 6e-g), suggesting that mancozeb selectively inhibits presynaptic GABA release in neurons. Similarly, 1-week oral administration of mancozeb in mice decreased both the frequency and amplitude of sIPSC (Fig. 6h-l), suggesting reduced astrocyte-mediated GABA release. Notably, this effect was abolished by coadministration of Ca^2+^ (Fig. 6h-l), indicating that Ca^2+^ rescues the mancozeb-induced reduction in GABAergic transmission.

**Fig. 6.**
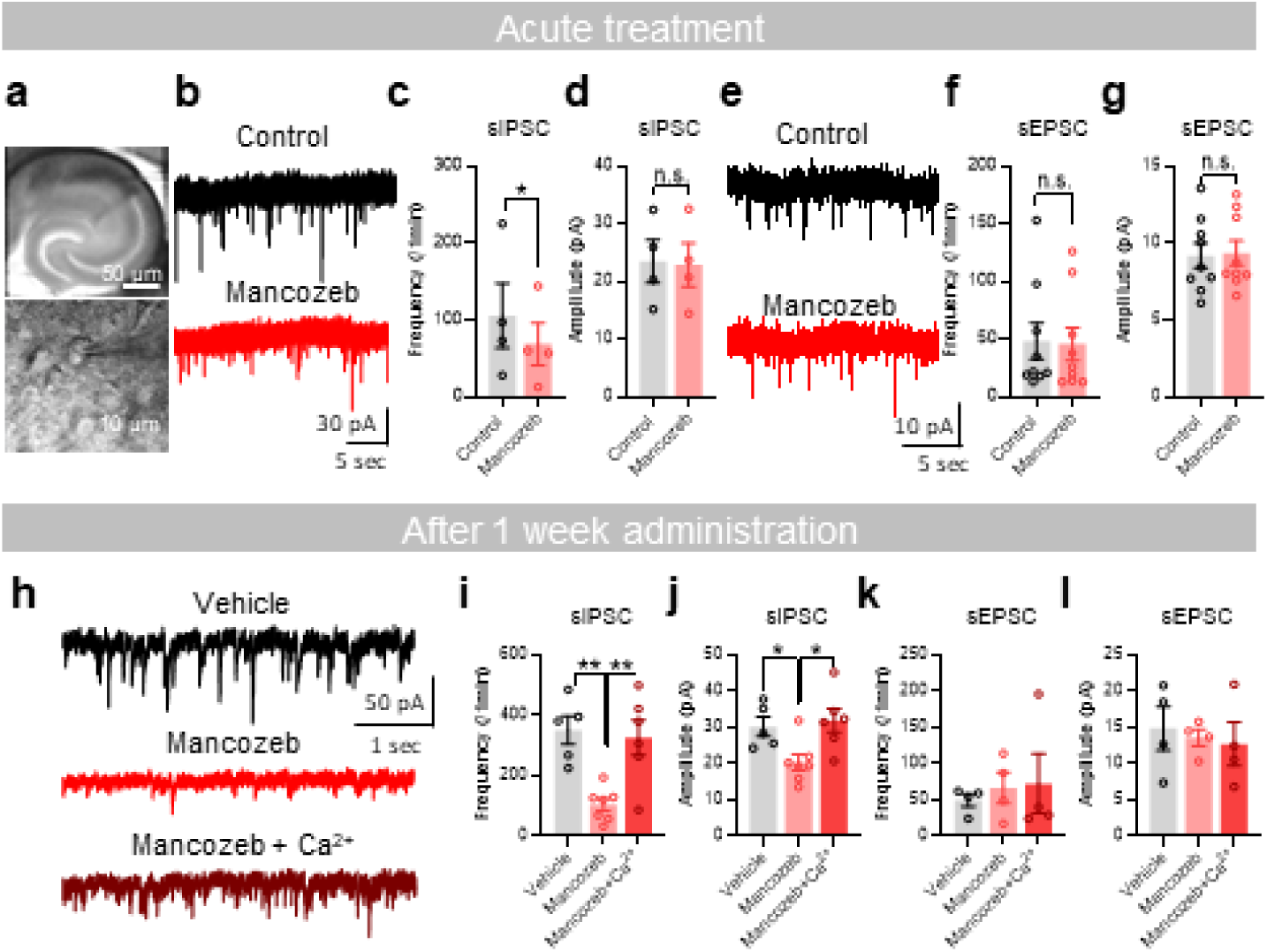
The supplementation of external Ca^2+^ rescues mancozeb-reduced sIPSC activity. **a** Bright-field images of the hippocampus. **b** Raw traces of sIPSC with pre- (black trace) and post-50 µM mancozeb (red trace). **c, d** Summary bar graph for the frequency **(c)** and the amplitude **(d)** of sIPSC. **e** Raw traces of sEPSC with pre- (black trace) and post-mancozeb (red trace). **f, g** Summary bar graph for the frequency **(f)** and the amplitude **(g)** of sEPSC. **h** Representative traces of sIPSC with vehicle (upper), mancozeb (middle), and mancozeb+Ca^2+^ (lower). **i, j** Summary bar graph for the frequency **(i)** and the amplitude **(j)** of sIPSC. **k, l** Summary bar graph for the frequency **(k)** and the amplitude **(l)** of sEPSC. Statistical significance was evaluated with one-tailed paired Student’s t-test (**c, d, f,** and **g,** *p < 0.05) and one-way ANOVA followed by Tukey’s test for multiple comparisons (**i** and **j,** *p < 0.05, **p < 0.01). Data are presented as mean ±SEM.

## 4. Discussion

Astrocytic GFAP atrophy in the hippocampus and entorhinal cortex has been strongly associated with psychiatric disorders such as schizophrenia and depression, despite the absence of any overt glial cell death [43–45]. However, the molecular mechanisms underlying this structural degeneration remain unclear. In our study, we demonstrated that mancozeb induced astrocytic atrophy, predominantly in the hippocampus and corpus callosum, accompanied by increased locomotor activity in the open-field test. Notably, treatment with SKF96365, an Orai1 channel inhibitor, similarly resulted in GFAP atrophy, suggesting a mechanistic link to SOCE. Mancozeb suppressed ER Ca²⁺ replenishment via Orai1/STIM1-mediated SOCE, as evidenced by the reduced amplitude and delayed latency of receptor-mediated Ca²⁺ transients. Furthermore, mancozeb-induced astrocytic atrophy selectively suppressed IPSC amplitude without affecting excitatory transmission, indicating disrupted inhibitory signaling. Based on these findings, we propose that impaired ER Ca²⁺ homeostasis due to SOCE dysfunction is a key cellular mechanism driving GFAP atrophy and selective inhibitory circuit imbalance.

Astrocyte atrophy has been attributed to multiple converging mechanisms, including mitochondrial dysfunction [46], impaired ER calcium homeostasis [47], and chronic inflammatory signaling [47]. These alterations compromise the astrocytic metabolic support, intercellular communication, and synaptic regulation, ultimately contributing to neural circuit instability and age-related neuropathology. The reduction in intracellular calcium (Ca²⁺) levels associated with the activation of the NO/cGMP signaling pathway induced by inflammatory cytokines may suppress GFAP expression and lead to structural atrophy of astrocytes [48]. Our findings demonstrate that mancozeb-induced astrocytic atrophy, coupled with pharmacological and gene silencing evidence of SOCE inhibition, provides the first mechanistic insight into how mancozeb impairs cytosolic Ca²⁺ replenishment from the ER to subsequently regulate intracellular GFAP expression.

Astrocytes preferentially interact with GABAergic interneurons through receptor-mediated calcium signaling and targeted gliotransmitter release, underscoring their essential role in modulating the inhibitory balance and neural circuit integration [49,50]. In line with these observations, our data demonstrated that mancozeb-induced suppression of ER Ca²⁺ levels in astrocytes, together with the inhibition of GABA release from inhibitory neurons, supports the notion that astrocytes specifically regulate inhibitory neuron activity. This disruption, accompanied by signs of astrocytic atrophy, may represent a key cellular mechanism underlying the imbalanced inhibitory control. In the intermediate region of the lateral septum, astrocytic Gq activation elicits a bidirectional shift in synaptic transmission, enhancing excitatory input while suppressing inhibition, thus underscoring the selective vulnerability of GABAergic neurons likely driven by astrocytes’ preferential connectivity and functional coupling with distinct neuronal subtypes [51]. The selective degeneration of inhibitory neurons is driven by astrocytic structural atrophy and impaired ER Ca²⁺ storage capacity. Our study is the first to elucidate glial dysfunction as a central mechanism underlying inhibitory circuit collapse. This degeneration likely disrupts gliotransmitter release as proper Ca²⁺ homeostasis is essential for vesicular signaling between astrocytes and neurons.

Astrocytic Ca²⁺ elevation in response to dopaminergic signaling modulates synaptic transmission via ATP/adenosine-mediated inhibition, thereby contributing to acute behavioral changes such as amphetamine-induced hyperlocomotion [52]. Astrocytic Ca²⁺ signaling in the hippocampus is sufficient to drive NMDA-dependent synaptic potentiation and facilitate task-specific memory enhancement, thus underscoring the critical role of glial calcium dynamics in neural plasticity and cognitive processing [53]. The influence of receptor-induced cytosolic calcium surges on cognition and memory in glial cells has been actively studied in recent years. However, this study is the first to propose that the disruption of calcium homeostasis in glia may contribute to atrophy.

Considering the established ADI for mancozeb at 0.023 mg/kg/day [54], our study employed sub-ADI dosing regimens (0.03 mg/kg/day for 1 week and 0.005 mg/kg/day for 3 weeks) to investigate the cumulative and temporal effects of low-level exposure on astrocytic calcium homeostasis and inhibitory circuit integrity. Notably, astrocytes exposed to 0.03 mg/kg/day mancozeb for 1 week exhibited aberrant calcium oscillation patterns and reduced buffering capacity, while prolonged exposure at 0.005 mg/kg/day over 3 weeks led to compromised inhibitory synaptic integrity. These alterations occurred in the absence of overt cytotoxicity, indicating that functional neurotoxicity may precede structural damage.

## 5. Conclusion

In conclusion, this study presents the first evidence that the fungicide mancozeb potently inhibits Ca^2+^ signaling and structural maintenance in astrocytes. Administration of mancozeb induced a decrease in GFAP expression, leading to structural damage in astrocytes. Mechanistically, we identified a mechanism demonstrating that mancozeb selectively inhibited Orai1-STIM1-mediated SOCE, and that this effect was abolished by Orai1 or STIM1 knockdown. This supports the fact that these channels are signaling targets of mancozeb. Notably, this inhibition led to a decrease in ER Ca^2+^ storage, as evidenced by the reduced Ca^2+^ response following P2Y1 activation. These effects in astrocytes ultimately reduced inhibitory synaptic transmission in neurons. Collectively, the results of this study suggest a novel astrocyte mechanism in which Mancozeb disrupts calcium ion signaling and consequently induces morphological abnormalities. This implies the possibility that Mancozeb contributes to the development of brain pathology through calcium ion signaling and morphological abnormalities.

## CRediT authorship contribution statement

**Ye-Ji Kim:** Conceptualization, Data curation, Formal analysis, Investigation, Methodology, Validation, Visualization, Writing – original draft, Writing – review and editing.

**Dong Ho Woo:** Conceptualization, Data curation, Funding acquisition, Project administration, Writing – original draft, Writing – review and editing.

## Acknowledgements

This work is supported by the National Research Council of Science & Technology (NST) grant by the Korea government (MSIT) (No. GTL24022-000); by Korea Institute of Marine Science & Technology Promotion(KIMST) funded by the Ministry of Oceans and Fisheries(RS-2025-02292973) to D.H.W. Jae Jung Lee performed western blotting for GFAP quantification and helped with animal experiments. The authors thank Dr. C. Justin Lee of the Institute of Basic Science (IBS, Daejeon, Korea) for providing shRNA constructs of Orai1, STIM-1, TRPA1, TRPC1, and Scrambles.

## Data availability

Data will be made available on request.

